# Ask1 and Akt act synergistically to promote ROS-dependent regeneration in *Drosophila*

**DOI:** 10.1101/451070

**Authors:** Paula Santabárbara-Ruiz, José Esteban-Collado, Lidia Pérez, Giacomo Viola, Marco Milán, Montserrat Corominas, Florenci Serras

## Abstract

The mechanism by which apoptotic cells release signals that induce undamaged neighbor cells to proliferate and regenerate missing parts remains elusive. Oxidative stress originated by dying or damaged cells can be propagated to neighboring cells, which then promote regeneration. We investigated the nature of the stress sensing mechanism by which neighboring cells are recruited. We found that *Drosophila* apoptosis signal-regulating kinase 1 (Ask1) senses reactive oxygen species (ROS) differently in stressed dying cells and unstressed neighboring cells and this differential sensing is pivotal for tissue repair. In undamaged cells, this activity is attenuated, but not abolished, by Akt1 phosphorylation, which thus acts as a survival signal that results in the tolerable levels of p38 and JNK necessary for regeneration. These observations demonstrate that the non-autonomous activation of the ROS-sensing mechanism by Ask1 and Akt1 in neighboring unstressed cells. Collectively, these results provide the basis for understanding the molecular mechanism of communication between dying and living cells that triggers regeneration.

**Author summary:** One of the early events that occur after tissue damage is oxidative stress production that signals to initiate wound healing and regeneration. Several signaling pathways, such as JNK and p38, respond to oxidative stress and are necessary for regeneration. We decided to explore the mechanism that links the oxidative stress and the activation of these pathways. We used epithelia of *Drosophila* to genetically direct cell death in specific zones of the tissue as means of experimentally controlled cell damage. We found that the Ask1 protein, which is sensitive to oxidative stress, is a key player in this scenario. Actually it acts as an intracellular sensor that upon damage activates those signaling pathways. However, high activity of Ask1 can be toxic for the cell. This is controlled by Akt, an enzyme dowstream the insulin pathway, with attenuates the activity of Ask1 to tolerable levels. In conclusion, Ask1 and Akt act synergistically to respond to the stress generated after tissue damage and drive regeneration. In other words, we found that the link between oxidative stress and nutrition is key for tissue regeneration.

## Introduction

Organisms are continuously exposed to a wide variety of environmental stressors that cause deterioration and cell death. Tissues overcome the effect of stressors by replacing or repairing damaged cells. Understanding the early signals that initiate the response to damage is an essential issue in regenerative biology. Compiling evidence supports that reactive oxygen species (ROS) fuel wound healing and oxygen-dependent redox-sensitive signaling processes involved in damage response [1,2].

Regeneration can be studied in *Drosophila* imaginal discs, which are epithelial sacs of the larvae, via genetic ablation of specific zones and then monitoring the mechanism of recovery by surviving cells [3]. Genetically induced apoptosis in the wing imaginal discs leads to production of ROS which propagate to surviving neighbors [4–6]. Although oxidative stress has been associated with several pathologies, low levels of ROS can be beneficial for signal transduction [7].

Jun-N Terminal kinase (JNK) and p38 are MAP kinases that respond to many stressors, including ROS, and foster tissue repair and regeneration in *Drosophila* [4,5,15–24,6,8–14]. Both signaling pathways control many cellular processes as disparate as cell proliferation and cell death. For example, ectopic activation of JNK induces apoptosis [25,26], but its inhibition results in lethality [27]. These disparities could be due to either different levels of activity or different mechanisms of activation.

Two different scenarios have been found after genetic ablation in the wing imaginal disc: (1) apoptotic cells show high concentration of ROS and high activity of JNK in the absence of p38 activity; and (2) the neighboring unstressed living cells show tolerable levels of ROS needed to trigger the low activity of JNK and the activation of p38 necessary for tissue regeneration [4]. A key question is how the balance between the beneficial or detrimental effects of ROS is controlled and, in particular, how ROS control JNK and p38 activity. A candidate molecule to perform this function is the MAPKKK Apoptosis signal-regulating kinase 1 (Ask1) which responds to various stresses by phosphorylation of JNK and p38 pathways [28–30]. Hence, in a reduced environment, thioredoxin (Trx) inhibits Ask1 kinase activity by directly binding to the N-terminal region of Ask1. Upon oxidative stress, the redox-sensitive cysteines of Trx become oxidized, resulting in the dissociation of Trx from Ask1. Consequently, Ask1 is oligomerized and its threonine-rich kinase domain is phosphorylated, inducing Ask1 activation [31,32]. Mammalian Ask1 is highly sensitive to oxidative stress and contributes substantially to JNK-dependent apoptosis. Nevertheless, recent studies have also revealed other functions of this kinase, including cell differentiation and survival [28,33].

Ask1-interacting proteins promote conformational changes that lead to the modulation of Ask1 activity and result in various cellular responses. For example, Akt1, a kinase activated by Pi3K pathway in response to insulin receptor activation, phosphorylates Ask1 and mitigates its activity *in vitro* [34].

In this work, we used wing imaginal discs, which encompass high regenerative capacity [3], to explore the link between ROS and regeneration. We found that Ask1 acts as a sensor of ROS upstream the JNK and p38 and that Akt1 is necessary for modulating Ask1 activity in undamaged regenerating cells. Together, our results indicate that oxidative stress generated in the damaged cells signals the neighboring undamaged cells to promote tissue repair.

## Results

The Gal4/UAS/Gal80^TS^ transactivation system is a key tool that allows us to activate, temporarily and in a spatially controlled manner, pro-apoptotic genes such as *reaper* (*rpr*) [9,10,18]. To determine if *Drosophila Ask1* is involved in regeneration, we first induced cell death in the wing disc using the wing-specific *sal^E/Pv^*-*Gal4* strain to activate *UAS-rpr* (henceforth *sal^E/Pv^>rpr*) in *Ask1* mutant backgrounds and then scored regeneration defects in adult wings. We found that in *Ask1^MB06487^* or *Ask1^MI02915^* heterozygous individuals, full regeneration of the wings dropped to 15% and 23%, respectively, in comparison to wild-type backgrounds (Fig. 1A). In addition, we used a double transactivation system to simultaneously express *rpr* in the *sal^E/Pv^* domain and the RNAi of *Ask1* in an adjacent compartment (Fig. 1B). The *UAS-Ask1^RNAi^* transgene was activated in the anterior compartment using the Gal4-UAS system (*ci-Gal4 UAS-Ask1^RNAi^*) and cell death was induced in the *sal^E/Pv^* domain using the Gal80-repressible transactivator system LHG, a modified form of the lexA lexO system (*sal^E/Pv^-LHG lexO-rpr*) (Fig. 1B) [4,35]. The resulting wings (*ci>Ask1^RNAi^ sal^E/Pv^>rpr*) lacked some veins or interveins and their size was reduced. In the absence of cell death (*sal^E/Pv^*> OFF in Fig.1B), *Ask1^RNAi^* wings were normal. Driving *UAS-Ask1^RNAi^* to a different compartment (dorsal) and inducing cell death in *sal^E/Pv^* (*ap>Ask1^RNAi^ sal^E/Pv^>rpr*) resulted in anomalies in 82% of wings. To evaluate whether these anomalies were due to impairment of the regenerative growth, we analyzed the mitotic index, calculated as the number of cells positive for the phosphorylated form of Histone 3 (P-H3) in the anterior compartment, where control and *Ask1^RNAi^* transgenes were expressed. We found an increase of mitosis in *sal^E/Pv^>rpr* discs, compared to the *sal^E/Pv^>GFP ci>RFP* or *sal^E/Pv^>GFP ci>Ask1RNAi* controls. However, this increase in the anterior compartment mitosis associated to damage, was blocked in *ci>Ask1^RNAi^* discs (Fig.1C), confirming that Ask1 reduction impairs regenerative growth.

**Figure 1.**
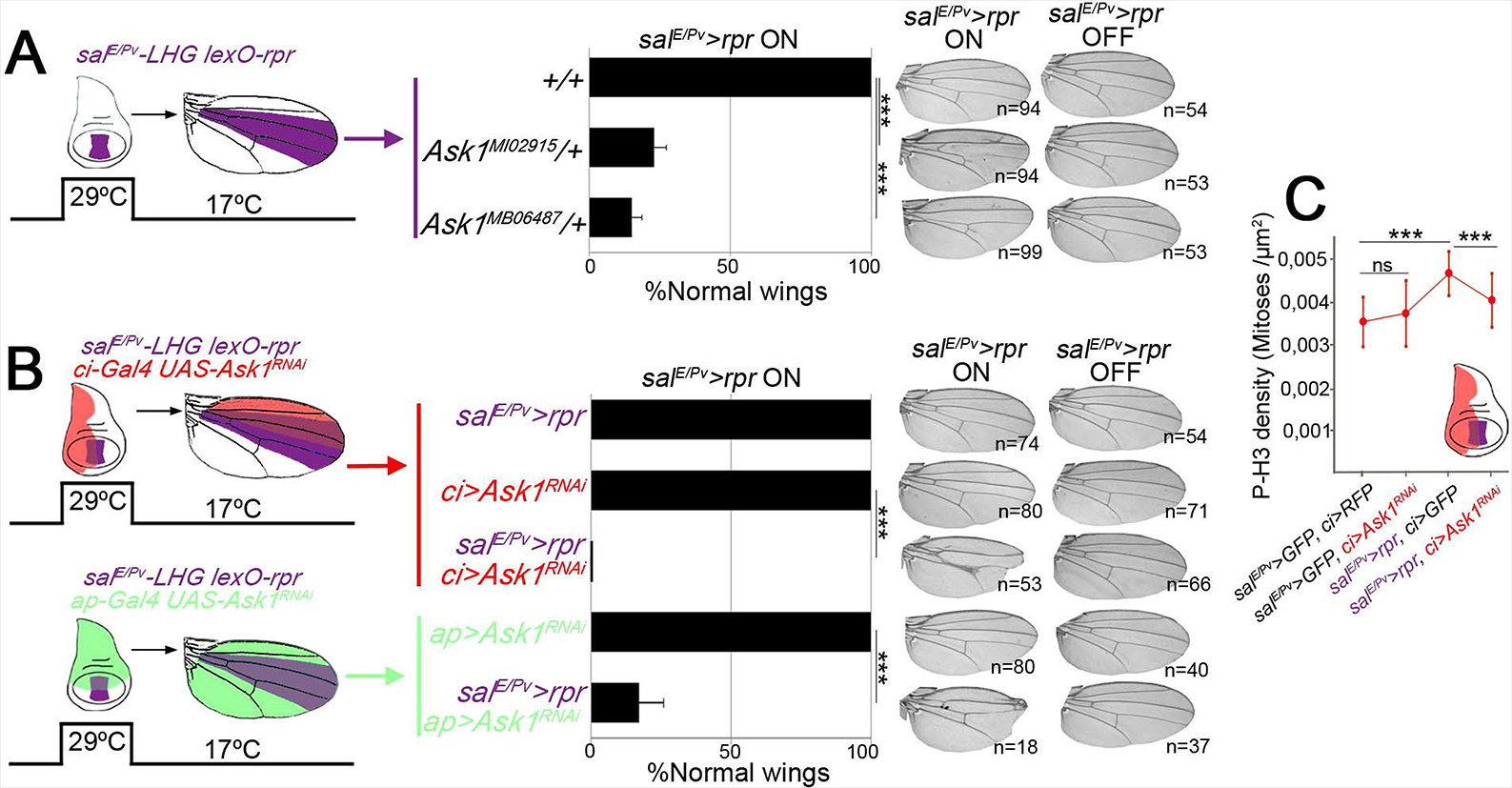
Ask1 is necessary for imaginal disc regeneration. (A) Percentage of fully regenerated wings (normal wings) in *Ask1* mutant backgrounds after genetic ablation. Left: Scheme of the zone ablated (purple) in wing disc and its corresponding region in the adult wing. Right: Examples of wings with full regeneration and incomplete regeneration. (B) Percentage of normal wings in controls (*rpr* or *Ask1^RNAi^* expression alone) and experimental samples (*rpr* and *Ask1^RNAi^* dual expression). Left: Double transgene activation scheme. Zone ablated using *sal^E/Pv^LHG lexO-rpr* (purple) in wing disc and its corresponding region in the adult wing; the zone of the *Ask1^RNAi^* transgene expression is indicated in red or green. Right: Examples of resulting wings. *Sal^E/Pv^>rpr* ON: indicates activated genetic ablation (11hours at 29°C);*sal^E/Pv^>rpr* OFF: examples of control wings without *rpr*-activation (kept at 17°C) for the genotypes indicated. (C) Phosphorylated H3 as indicator of mitotic cells per area in discs with the dual transactivation system using *rpr* and *Ask1^RNAi^* for the genotypes indicated (n=30 each genotype). Error bars in (A,B) show standard error of sample proportion and in (C) standard deviation. ***P<0.001.

The *Drosophila Ask1* locus encodes two peptides of different lengths, Ask1-RB and Ask1-RC, with predicted molecular weights of 136.5 kDa and 155.5 kDa, respectively. Both have a protein kinase-like domain that contains a highly conserved core of threonines conferring functionality on the protein [28]. The longer Ask1-RC isoform contains a conserved domain of unknown function DUF4071 (Fig. 2A) in which, in mammals, phosphorylation of the Ser83 results in attenuation of Ask1 activity in vitro [34]. Active Ask1 can be traced with antibodies against the phosphorylated threonine residues of the highly conserved kinase domain (henceforth P-Thr) and the attenuated form with specific antibodies against phosphorylated Ser83 (P-Ser83). We first tested both antibodies in wild-type wing imaginal discs. We detected low levels of P-Thr all over the disc including some transient high activity during mitosis (Fig. 2B and S1A Fig.), as occurs in mammalian cells [36]. We also found low levels of P-Ser83 in wild-type control discs (Fig. 2C). Both the general low and the high mitotic-associated activity of P-Thr as well as the low endogenous P-Ser83 levels, were abolished in *Ask1* mutant backgrounds (S1B, C Fig.).

**Figure 2.**
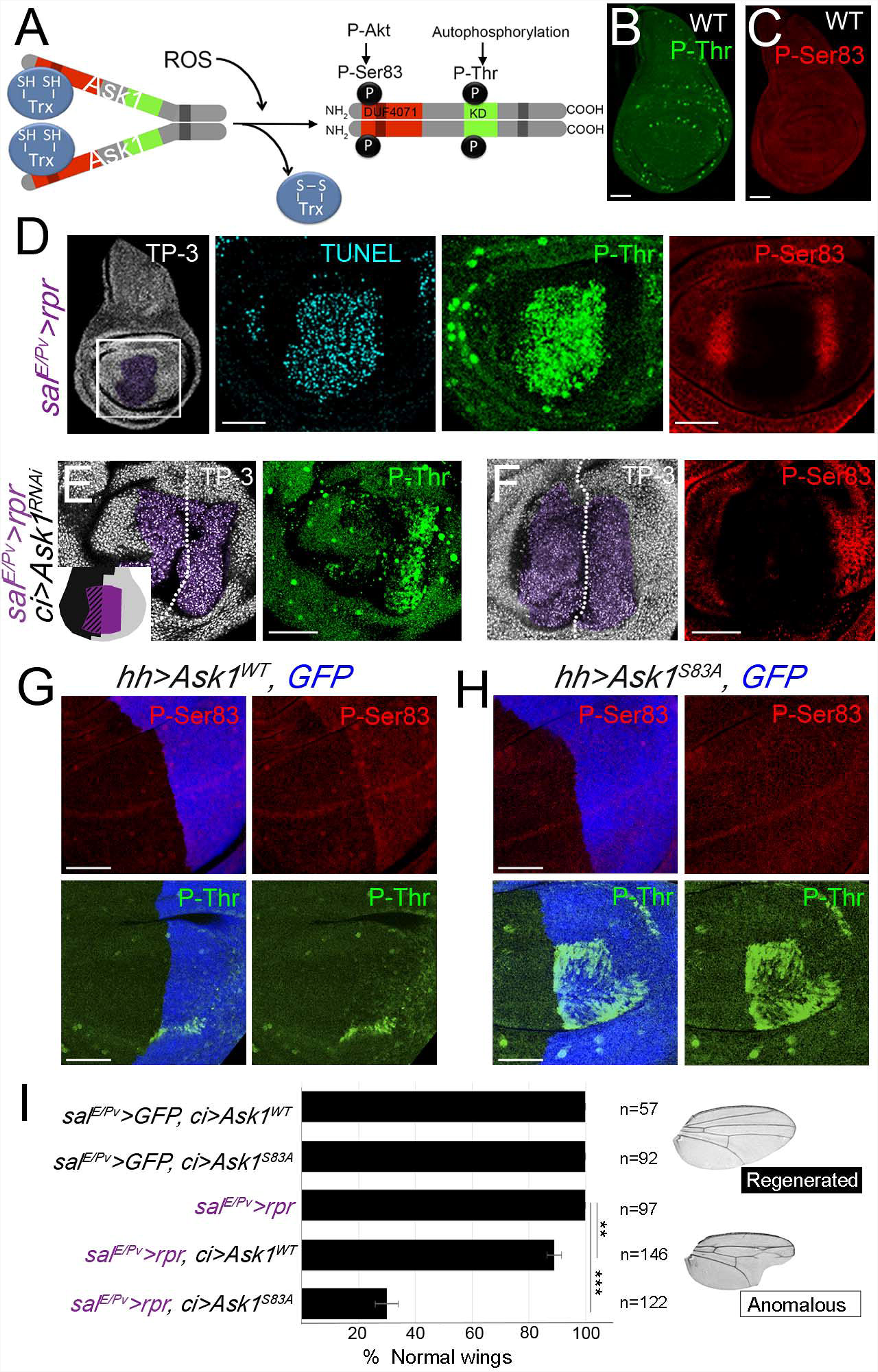
Ask1 is activated upon induction of apoptosis. (A) Structural features of Ask1-RC isoform. It possesses a phosphokinase domain (KD) with a highly conserved core of threonines (in green). In the N-terminal domain, Ask1-RC shows the DUF4071 domain (in red). In the C-terminal domain, there is a coiled coil domain (dark grey), which allows interactions between Ask1 monomers to form oligomers [28]. Phosphorylation sites are shown. Activation of Ask1 is ROS-dependent. Inactive Ask1 is constituted by association of Ask1 monomers through the coiled coil domain and the reduced form of thioredoxin (Trx-SH) in the N-terminal domain. Upon oxidation, Trx-SS dissociates, which allows Ask1 to autophosphorylate in the phosphokinase domain, leading to an active Ask1. Active Akt attenuates Ask1 activity by phosphorylating the Ser83 residue *in vitro*[28,34]. (B-C) Characterization of mammalian P-ASK1 antibodies in *Drosophila* discs. Endogenous levels of P-Thr (B) and P-Ser83 (C) in wild-type (WT) discs. P-Thr is found in all cells, but increases in mitotic cells. P-Ser83 is also found in all cells of the disc epithelium. (D) Ask1 activity in *sal^E/Pv^>rpr* genetic ablation. *Sal^E/Pv^>rpr* disc indicating the area imaged (white square); TP-3 (TO-PRO-3) nuclei; purple area: the *sal^E/Pv^>rpr* ablated zone. Disc stained with Ask1 P-Thr, detected in apoptotic cells (TUNEL assay, n=21). Disc stained with Ask1 P-Ser83, only found in living cells near the apoptotic zone (n=35). (E-F) After cell death (*sal^E/Pv^>rpr*) and simultaneous inhibition of *Ask1* in the anterior compartment (*ci>Ask1^RNAi^*; black zone in the inset cartoon), P-Thr (E) and P-Ser83 (F) were specifically inhibited in this area. White dotted line indicates the AP boundary; anterior to the left, posterior to the right. The inset in (E) depicts the *Ask1^RNAi^* area (black) and the *sal^E/Pv^>rpr* ablated zone (purple). (G) Ectopic expression of the *Ask1^WT^* form in the posterior compartment (blue) leads to an increase in P-Ser83 and a weak increment of P-Thr. (H) Ectopic expression of the truncated form *Ask1^S83A^* does not raise levels of P-Ser83, but does increase PThr. Scale bars 50µm. (I) Percentage of regenerated wings (normal wings) in controls (*rpr*, *Ask1^WT^or Ask1^S83A^* expression alone) and experimental samples (*rpr* and *Ask1^WT^or Ask1^S83A^* dual expression). Right: Examples of a wing with normal and anomalous regeneration (*sal^E/Pv^>rpr, ci> Ask1S83A*). Error bars show standard error of sample proportion. **P<0.01 ***P<0.001.

However, upon apoptosis, high levels of P-Thr were localized in dying cells (Fig. 2D) and absent or, in some cases, very weakly incremented above the basal levels in nearby living cells. This increment of P-Thr in apoptotic cells was abolished after *Ask1^RNAi^* expression (Fig. 2E). In contrast, P-Ser83 accumulated in living cells adjoining the apoptotic zone and was absent in apoptotic cells. The increase in P-Ser83 varied from discs with strong accumulation near the dying domain to those with an extended increase in the whole wing pouch (Fig. 2D and S2A Fig.). Similar results were obtained after killing cells with a different pro-apoptotic gene (*sal^E/Pv^>hid*) (S2B Fig.). In the presence of apoptosis and blocking *Ask1*, the P-Ser83 increment in living cells was inhibited (Fig. 2F). P-Ser83 was also found to be elevated at the wound edges after physical injury (S2C Fig.). Together, these observations indicate that neighboring cells respond to damage by phosphorylation of Ask1 Ser83.

In contrast to the high activity of Ask1 in dying cells (high P-Thr), the presence of P-Ser83 in living cells could be indicative of tolerable levels of Ask1 achieved by attenuation of P-Thr activity. To test this hypothesis, we mutated *Ask1* at serine 83 to alanine and cloned it into a UAS vector (*UAS-Ask1^S83A^*). In parallel, we also cloned a wild-type form of Ask1 (*UAS-Ask1^WT^*). The ectopic expression of *Ask1^WT^* resulted in an increase of P-Ser83 in addition to low levels of P-Thr (Fig. 2G). Interestingly, the expression of *Ask1^S83A^*, which did not show a rise in P-Ser83, resulted in high levels of P-Thr (Fig. 2H). This observation concurs with the P-Ser83 residue as responsible for the attenuation of P-Thr.

We also analyzed whether the Ser83 residue is key for damage response. Using the double transactivation system, we found that the expression of *UAS-Ask1^WT^* or *UAS-Ask1^S83A^* transgenes caused no effects in the absence of stressed tissue (Fig. 2I). Upon apoptosis, expression of *UAS-Ask1^WT^* resulted in normal regeneration in 89% of wings. In contrast, expression of *UAS-Ask1^S83A^* led to a fall to only 29% of individuals being capable of regenerating and the rest showed strong effects on veins and interveins, as well as the appearance of notches, which together are indicative of disrupted regeneration (Fig. 2I). The suppression of the ability to regenerate by *UAS-Ask1^S83A^*, which may act as a dominant negative allele, is likely due to the lack of non-autonomous P-Ser83 increment in these cells.

Next, we decided to identify the upstream signal responsible for Ser83 phosphorylation. In mammalian cells, phosphorylation of Ask1 at Ser83 is driven by the serine-threonine Akt kinase, a key signaling molecule in the insulin pathway [34]. We wondered whether the *Drosophila* Akt1 as well as the Pi3K92E kinase (the *Drosophila* Pi3K kinase also known as dp110), were required for P-Ser83 activity. We first found that the active phosphorylated form of Akt (P-Akt) was increased in the living tissue and decreased or was absent in the dying zone (Fig 3A-C). This suggests that Akt1 functions in surviving cells. To test whether the Akt1 phosphorylated the endogenous Akt1 Ser83, we used an RNAi for *Akt1* and found that P-Ser83 was absent (Fig. 3D, E). In addition, ectopic activation of Akt1 (*UAS-myr-Akt1.S*) [37] resulted in an increase in the levels of P-Ser83 (Fig.3F, G).

**Figure 3.**
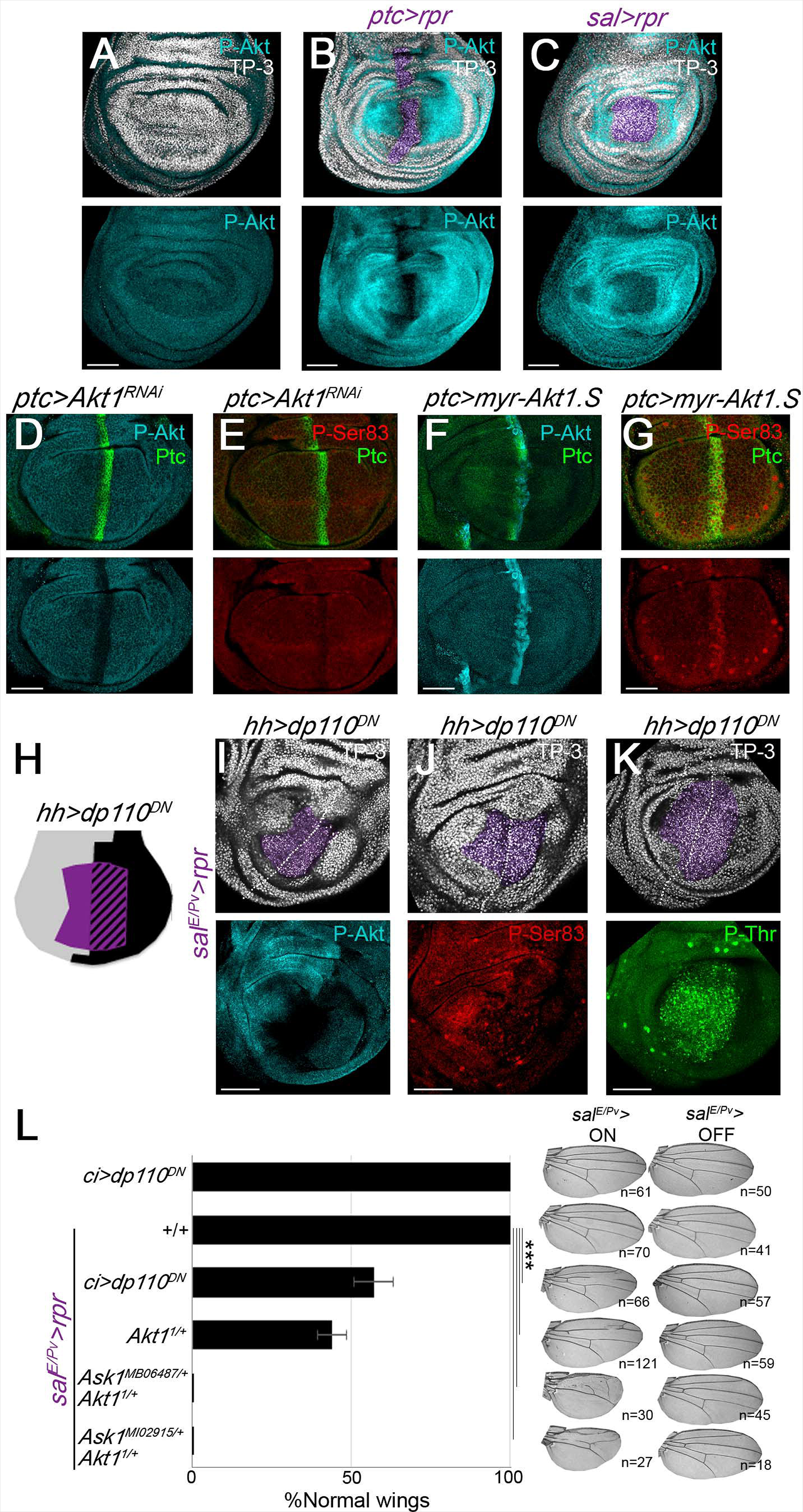
Phosphorylation of *Drosophila* Ask1 Ser83 depends on the Pi3K/Akt1 pathway. (A-C) P-Akt staining in wild-type discs (A), and after *ptc>rpr*(B), and *sal>rpr* induction (C). Akt1 is activated in living cells surrounding the apoptotic cells. Apoptotic zone of pyknotic nuclei, in purple. (D,E) RNA interference of *Akt1* in the *ptc* stripe of cells (*ptc>Akt1^RNAi^*) leads to a decrease of P-Akt and Ask1 P-Ser83 basal levels. (F,G) Ectopic expression of a constitutively active form of *Akt1*(*ptc>myr-Akt1.S*) promotes PAkt and Ask1 P-Ser83. (H) Design of the experiments in I,J and K with *sal^E/Pv^>rpr* induction (purple) and blocking Pi3K (*hh>dp110^DN^*) in the posterior compartment (black). Staining with anti-P-Akt (I), with anti-Ask1 P-Ser83 (J) with Ask1 P-Thr (K). Note that both P-Akt and P-Ser83 decrease in the posterior compartment but not PThr. Dotted white line indicates anterior-posterior boundary. TP-3; TO-PRO-3 nuclear staining. Scale bars 50µm. (L) Left: Percentage of fully regenerated wings after genetic ablation and Pi3K pathway inhibition alone or in combination with *Ask1* mutant backgrounds (genotypes indicated). Right: *sal^E/Pv^>* ON, examples of wings with full regeneration (controls *sal^E/Pv^>rpr, ci>dp110DN*) and anomalous regeneration (genotypes indicated); *sal^E/Pv^>* OFF wings: examples of control wings in the absence of *rpr*-ablation (maintained at 17°C). Error bars indicate standard error of sample proportion. ***P<0.001.

We also studied the role of P-Akt on Ask1 in the context of regeneration. We found that the dominant negative form of *Pi3K92E* (*dp110^DN^*) blocked the accumulation of P-Akt and P-Ser83 induced after genetic ablation (Fig. 3H-J), but not the P-Thr in dying cells (Fig. 3K and S3 Fig.). We also scored the effects on wing regeneration after blocking the Akt pathway. We observed impaired regeneration when the dominant negative form of *Pi3K92E* was expressed in the anterior compartment (*ci>dp110^DN^*) as well as in the *Akt1^1^* heterozygous background, an allele that encodes a catalytically inactive protein [38] (Fig. 3L). Moreover, regeneration was severely affected in double heterozygous flies containing *Akt1^1^* and *Ask1^MB06487^* or *Ask1^MI02915^* alleles (Fig. 3L). Neither the dominant negative form of *Pi3K92E* nor the allelic combination *Akt1^1^* and *Ask1^MB06487^* or *Ask1^MI02915^* affected wing development in the absence of cell death (*sal^E/Pv^>* OFF wings in Fig. 3L). These results further support the notion that Pi3K92E/Akt1 and Ask1 genetically interact and that their activity is key in driving regeneration.

Next, to test whether Ask1 activation is ROS-dependent, we fed *sal^E/Pv^>rpr* larvae with food supplemented with N-acetyl cysteine (NAC), a potent non-enzymatic scavenger that decreases ROS production, and examined Ask1 phosphorylation (Fig. 4A). After induction of cell death, we found a significant decrease in both P-Thr and PSer83 in discs of the NAC-fed larvae (Fig. 4B,C). In addition, we fed larvae with H_2_O_2-_supplemented food and observed a significant increase in both P-Thr and P-Ser83 levels (Fig. 4D-F). This oxidative stress-induced increase was partially blocked in the hypomorphic *Ask1^MB06487^* homozygous mutant (Fig. 4E, F). It is known that in mammalian cells Ask1 can also be activated by endoplasmic reticulum (ER) stress [39]. Feeding larvae with tunicamycin, an inhibitor of N-glycosylation in the ER that induces ER stress, led to an increase in Ask1 activation, which was also blocked in *Ask1^MB06487^* mutant discs (Fig. 4E, F).

**Figure 4.**
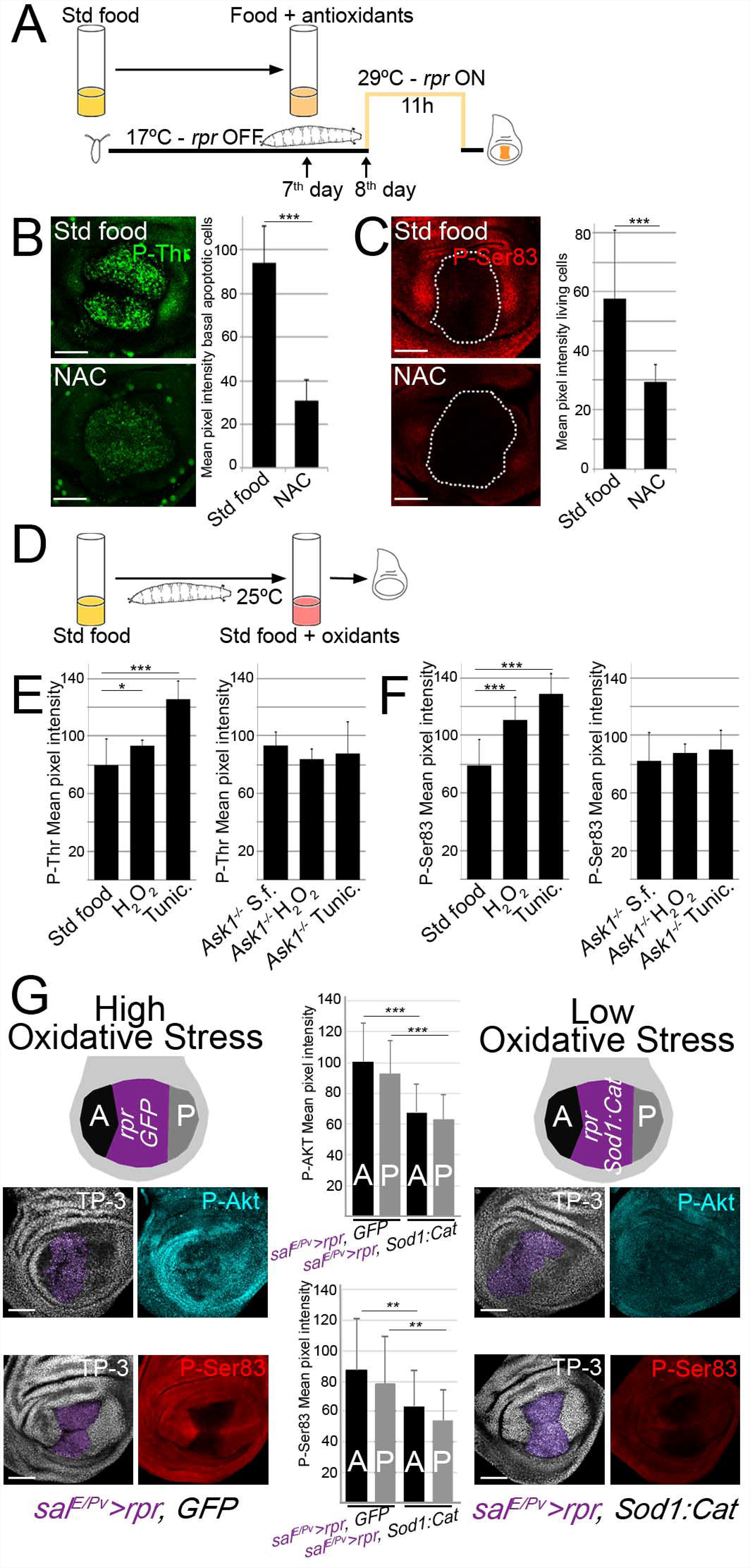
Ask1 is activated by ROS. (A) Design of experiments after feeding larvae with the NAC antioxidant. (B) Examples of Ask1 P-Thr after *sal^E/Pv^>rpr* induction in standard or NAC-supplemented food. Mean pixel intensities of P-Thr fluorescent labeling in the apoptotic zone of *sal^E/Pv^>rpr* discs from larvae fed with a standard (dying cells 94.01±17.04; S.D., n=11) or NAC-supplemented (dying cells 31.07±9.38; S.D. n=10) diet. (C) Examples of Ask1 P-Ser83 after *sal^E/Pv^>rpr* induction with standard or NAC-supplemented food. White dotted line indicates the *sal^E/Pv^>rpr* dying domain. Mean pixel intensities of Ask1 P-Ser83 fluorescent labeling in the neighboring cells near the apoptotic zone of *sal^E/Pv^>rpr* discs from larvae fed with a standard (living cells 57.56±23.27; S.D., n=20) or NAC-supplemented (living cells 29.54±5.92; S.D., n=18) diet. (D) Design of experiments after enhancing oxidative stress. (E) Left: P-Thr mean pixel intensity of wild-type wing discs from larvae fed with standard (79.36±18.65; S.D., n=17), H_2_O_2_-(93.37±3.87; S.D., n=7) and tunicamycin-supplemented (125.49±12.51; S.D., n=7) food. Right: P-Thr mean pixel intensity of *Ask1^MB06487^* homozygous mutant discs from larvae fed with standard (92.92±9.75; S.D., n=6), H_2_O_2_-(83.66±7.11; S.D. n=11) and tunicamycin-supplemented (87.99±21.38; S.D., n=7) food. (F) Left: P-Ser83 mean pixel intensity of wild-type wing discs from larvae fed with standard (79.14±17.69; S.D., n=19), H_2_O_2_-(110.89±15.43; S.D., n=14) and tunicamycin-supplemented (128.35±14.42; S.D., n=5) food. Right: P-Ser83 mean pixel intensity of *Ask1^MB06487^* homozygous mutant discs from larvae fed with standard (81.83±20.25; S.D., n=12), H_2_O_2_-(87.51±6.40; S.D., n=6) and tunicamycin-supplemented (90.27±13.26; S.D.; n=4) food. (G) P-Akt (top) and P-Ser (bottom) mean pixel intensity from discs with *rpr* activation (*sal^E/Pv^>rpr, GFP;* high stress) and from discs with simultaneous *Sod1:Cat* and *rpr* activation (*sal^E/Pv^>rpr, Sod1:Cat;* low stress). Purple area shows where transgenes were activated. A and P are the zones where intensity was measured. PAkt mean pixel intensity for *sal^E/Pv^>rpr, GFP* was 100.61±25,14, S.D. in A and 92.64±21.38, S.D. in P; and for *sal^E/Pv^>rpr, Sod1:Cat* was 68,04±18,16, S.D. in A and 63.14±15.66, S.D. in P. P-Ser83 mean pixel intensity for *sal^E/Pv^>rpr, GFP* was 86.89±34.49, S.D. in A and 78.46±31.06, S.D. in P; and for *sal^E/Pv^>rpr, Sod1:Cat* was 63.25±23.82, S.D. in A and 54.32±19.92, S.D. in P. N=23 discs for each genotype. *P<0.05. **P<0.01, ***P<0.001. Error bars indicate standard deviation. TP-3: TO-PRO-3 nuclear staining. Scale bars: 50µm.

Furthermore, we wondered whether Akt1 activation in the unstressed living cells is targeted by ROS from the dying cells. To examine this hypothesis, we enzymatically blocked ROS production using ectopic expression of the ROS scavengers *Superoxide dismutase 1* and *Catalase* (*Sod1:Cat*) in the *rpr* ablated zone. Remarkably, we found that activation of P-Akt was reduced when *Sod1:Cat* was co-expressed with *rpr* in the same zone (Fig. 4G). Accordingly, we found that the increment of P-Ser83 in neighboring cells was blocked when *Sod1:Cat* was co-expressed with *rpr* (Fig. 4G). These results demonstrate that the oxidative stress generated from the dying cells targets the Pi3K/Akt1 and Ask1 in the neighboring undamaged tissue.

We have shown that the Ask1 P-Ser83 attenuating signal is essential for regeneration, and that this signal depends on ROS and Akt. We next hypothesized that this P-Ser83-mediated attenuation does not silence Ask1, but instead maintains the low levels of Ask1 activity necessary for regeneration. Therefore we analyzed the expression of its known targets, the stress-activated MAP kinases JNK and p38 [40,41], both required for ROS-dependent regeneration [4–6]. We blocked *Ask1* in the anterior compartment and used the posterior as an internal control for the same disc (*ci>Ask1^RNAi^ sal^E/Pv^>rpr*) and found most phosphorylated p38 (P-p38), as an indicator of p38 activity, in the posterior compartment (Fig. 5A, B). The effects of Ask1 on JNK activity were monitored by matrix metalloproteinase 1 (Mmp1) expression as a read-out of the pathway [42]. After cell death, Mmp1 was found in both compartments, but in *ci>Ask1^RNAi^ sal^E/Pv^>rpr* discs Mmp1 levels dropped in the anterior (Fig. 5C, D). We also analyzed whether *Ask1* was required for activation of p38 and JNK after physical damage. We made two incisions in the same disc, one in the *UAS-Ask1^RNAi^* compartment and another into the control compartment. We found that P-p38 localized at the wound edges in the control compartment, whereas the levels of P-p38 were reduced at the wound edges of the *Ask1^RNAi^* compartment (Fig. 5E, G). Likewise, cut *Ask1^RNAi^* discs resulted in less Mmp1 than in the control compartment (Fig. 5F, H).

**Figure 5.**
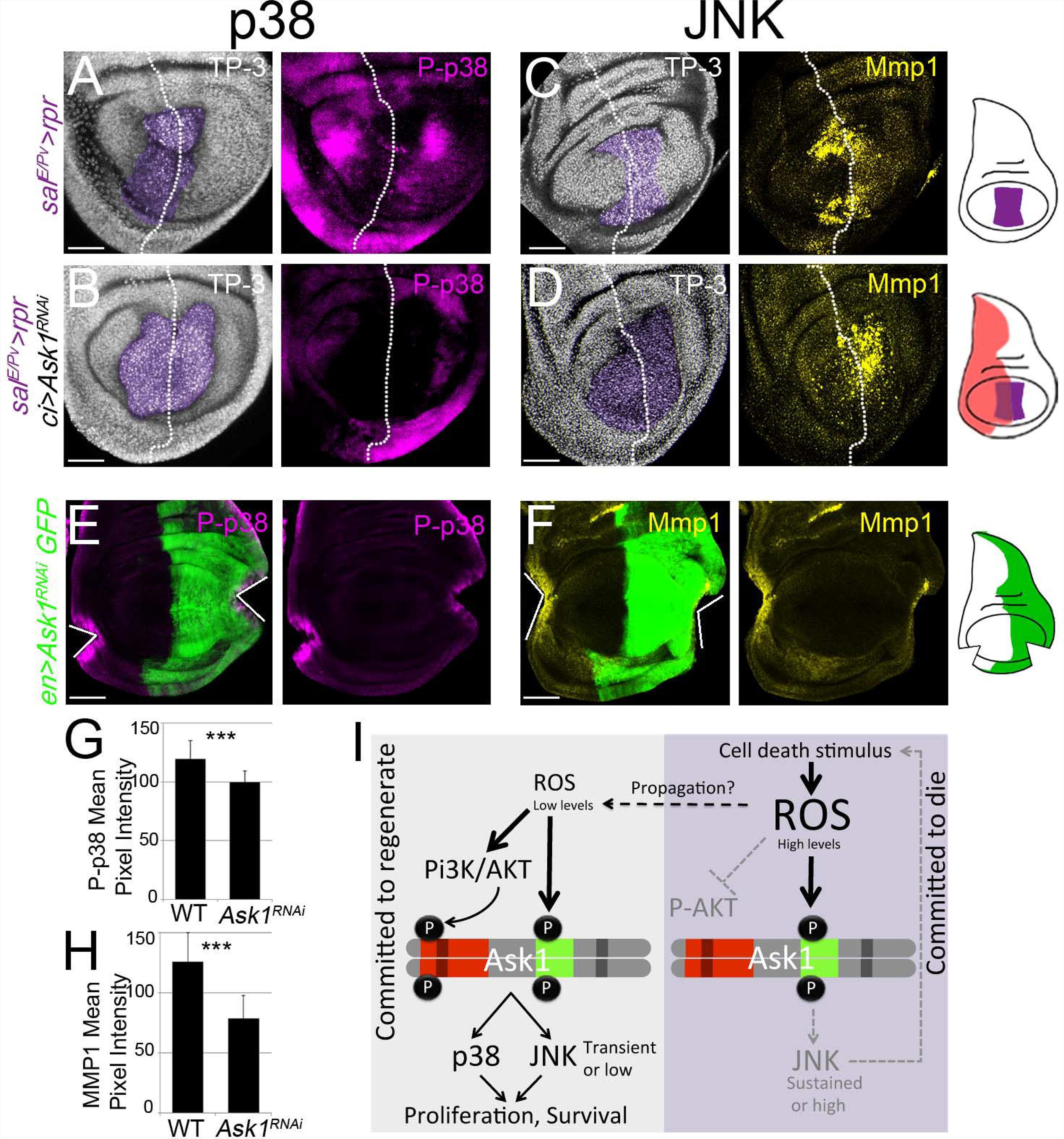
Ask1 promotes JNK and p38 signaling. (A) P-p38 after *sal^E/Pv^>rpr* induction. P38 is phosphorylated in living cells around the dying domain (n=14). (B) Inhibition of *Ask1* in the anterior compartment (*sal^E/Pv^>rpr ci>Ask1^RNAi^*) blocks induced P-p38 (n=7). (C) Mmp1 after *sal^E/Pv^>rpr* induction. Mmp1 expression is induced in dying and living cells (n=16). (D) Inhibition of *Ask1* in the anterior compartment (*sal^E/Pv^>rpr ci>Ask1^RNAi^*) blocks Mmp1 expression (n=14). TP-3: TO-PRO-3 nuclear staining. Purple: *sal^E/Pv^>rpr* ablated zone. White dotted line divides the discs into anterior (*Ask1* inhibition; to the left) and posterior (wild-type; to the right) compartments. (E-H) RNA interference of *Ask1* inhibits P-p38 and Mmp1 expression after physical injury. The *UAS-Ask1^RNAi^* was activated in the posterior compartment together with *UAS-GFP*(green). Two injuries were inflicted with tungsten needles in Schneider’s medium, one in the anterior and one in the posterior compartment. (E) For P-p38 staining, discs were immediately fixed after physical injury. (F) For Mmp1 staining, discs were first cut and cultured for 5 hours before fixation and processing. White lines indicate the wound site. Scale bar: 50µm. (G) Quantification of P-p38 activation at the wound site (n=10 discs each genotype). (H) Quantification of Mmp1 at the wound site (n=8 discs each genotype). Error bars indicate standard deviation. ***P<0.001. (I) Model proposed for the onset of regeneration. Low levels of ROS trigger regeneration through Ask1 activation, which is attenuated by the Pi3K/Akt pathway, leading to moderate activation of JNK and p38. In dead cells, high ROS levels promote high Ask1 and high JNK activity, in the absence of Akt1-driven attenuation.

## Discussion

We have shown here that Ask1 acts as a sensor of ROS after damage, and that synergizes with Pi3K/Akt1 to phosphorylate the Ask1 P-Ser83, a key phosphorylation event to initiate regeneration. In addition, we demonstrated that the activation of the Pi3K/Akt1 and Ask1 in undamaged cells is originated by ROS produced by the damaged tissue.

An essential question to understand regeneration is how damaged cells communicate to the nearby surviving cells to initiate regeneration. One of the most classical and determining studies on epithelial regeneration was the discovery that massive cell death induced with ionizing radiation in *Drosophila* larvae results in proliferation of the unharmed neighbors to compensate the lost [43]. Since then, compensatory proliferation, a cellular response linked to regeneration, has been considered as a result of signals released from apoptotic cells [44]. We propose here that the oxidative stress generated by damaged or apoptotic cells signals undamaged tissue to ignite the Ask1/Akt1 machinery that will culminate with repair and compensatory proliferation.

ROS acting in signaling after wounding has been subject of extensive research in various organisms and tissues [4,5,52–56,6,45–51]. Here, we have uncovered two scenarios of Ask1 activity: high activity in apoptotic cells, with high levels of P-Thr; and low activity in undamaged cells, where P-Ser83 prevails over P-Thr. This fits with the high levels of ROS and JNK in dying cells and the low levels of ROS, low JNK and P-p38 in undamaged cells reported previously [4]. Together, our observations address the question of how JNK signaling selectively fosters apoptosis or proliferation, and how this signal is related to high or low ROS levels (Fig. 5I). The same applies to p38, which is only activated in unharmed cells when neighbors enter apoptosis or are damaged [4,57]. Our study demonstrates that Ask1 operates in an Akt1-dependent manner in living cells and in an Akt1-independent manner in dying cells (Fig. 5I). In addition, it is conceivable that Akt1-mediated attenuation delivers either low or transient levels of Ask1 activity to stimulate P-p38 and low JNK levels necessary for tissue regeneration. In contrast, apoptotic cells induce high or sustained levels of JNK, either because of the amplification loop of apoptosis triggered by JNK [58] or because high levels of ROS result in high Ask1 P-Thr in the absence of Pi3K/Akt1 attenuation. We conclude that oxidative stress and the absence or presence of Pi3K/Akt1 survival signals are integrated in Ask1 to control the selective activity of JNK and p38, and in turn regenerative growth.

## Materials and Methods

### *Drosophila* strains

The *Drosophila melanogaster* strains *sal^E/Pv^-Gal4*, *tubGal80^TS^*, *UAS-rpr*, *sal^E/Pv^-LHG* and *lexO-rpr*, are previously described elsewhere [4]. *Ask1^MB06487^* [59] *Ask1^MI02915^*[60], *Akt1^1^* [38], *ptc-Gal4*, *sal-Gal4, ci-Gal4, ap-Gal4*, *en-Gal4*, *hh-Gal4*, *UAS-hid*, *UAS-GFP, UAS-RFP, dpp-Gal4, hh-Gal4, UAS-dp110^DN^* (D954A), *Df(3R)BSC636* (25726), *UAS-Sod.A (Sod1), UAS-Cat.A,* and *UAS-Ask1^RNAi^* (35331) were obtained from the Bloomington Stock Center. *UAS-Akt^RNAi^* (2902) was taken from the Vienna Drosophila Resource Center (VDRC). We used the *UAS-myr-Akt1.S,* a constitutively activated membrane-anchored form of *Akt1*[37] Canton S and *w^118^* were used as controls.

The *Ask1^MB06487^* mutant is viable in homozygosis and shows transcription after a RT-PCR (results not shown). The *Ask1^MI02915^* mutant is lethal in homozygosis as well as over the *Df(3R)BSC636.*

### Generation of *UAS-Ask1^WT^* and *UAS-Ask1^S83A^* flies

pUASt-attb_Ask1^WT^ was constructed by cutting Ask1 cDNA EcoRI /BamHI from DGRC clone FI02066 and cloning it into pUASt-attb.

pUASt-attb_Ask1^S83A^. Two serines in the 75 and 83 residues in the DUF4071 domain showed putative non-canonical AKT phosphorylation sites (ILTQQRPL<u>S</u>YHYHLGVRE<u>S</u>F). Both residues are spatially exposed similarly to human Ser83 of ASK1, which makes them accessible to kinases. As the AKT kinase phosphorylates the Ser83 of human ASK1 and attenuates the activity of ASK1 *in vitro* (Kim et al. 2001), we decided to mutate this residue and clone it into a UAS vector. pUASt-attb_Ask1 S83A was constructed by mutating serine 83 to alanine by PCR using oligos Ask1Mut-Fwd and Ask1S83A-Rev for partial PCR1 and Ask1S83A-Fwd and Ask1Mut-Rev for partial PCR2. Complete PCR was performed using the two partial PCRs as templates with oligos Ask1Mut-Fwd and Ask1Mut-Rev. The complete PCR was then cut with EcoRI /PflMI and cloned into FI02066-cut EcoRI/PflMI. The mutation introduced a new StuI site that was used to check the mutated clones. Mutated Ask1S83A cDNA was then cut from EcoRI/BamHI and cloned into pUASt-attb. Both clones were injected by standard procedures in line zh-86Fb-attP and transgenic lines were selected.

Ask1Mut-Fwd: AAT ACA AGA AGA GAA CTC TGA ATA CGG AAT

Ask1Mut-rev: CGG CGG TGT GGT TTT GTG CAC AAA CCG ATC

Ask1S83A-Fwd: CGT TAG GGA GGC CTT CGG GAT GAA GGA GA

Ask1S83A-Rev: CGG CGG TGT GGT TTT GTG CAC AAA CCG ATC

### Genetic ablation and dual Gal4/LexA transactivation system

Cell death was genetically induced as previously described [18,19]. We used the *sal^E/Pv^-Gal4* as a driver, which consists of *spalt* wing enhancer with expression confined to the wing [61] to score adult wing parameters. The UAS line used to promote cell death was *UAS-rpr* or *UAS-hid*, controlled by the thermo-sensitive Gal4 repressor *tubGal80^TS^*. We also used the *sal^E/Pv^-LHG* and *LexO-rpr* strains [4] for genetic ablation, utilizing the same design as for Gal4/UAS.

Embryos were kept at 17°C until the 8th day/192 hours after egg laying to prevent *rpr* expression. They were subsequently moved to 29°C for 11 hours and then back to 17°C to allow tissue to regenerate. Two types of controls were always treated in parallel; individuals without *rpr* expression (*UAS-GFP*, moved to 29°C for 11 hours) and individuals kept continuously at 17°C to avoid any transgene activation

In dual transactivation experiments, we used *sal^E/Pv^-LHG LexO-rpr* to ablate the *sal^E/Pv^* domain (abridged as *sal^E/Pv^>rpr*), whereas Gal4 was used to express different transgenes under the control of Gal4 drivers (*ci-Gal4* for anterior compartment; *ap-Gal4* for dorsal compartment and *hh-Gal4* for posterior compartment).

### Test for regenerated adult wings and statistics

To test the capacity of different genetic backgrounds to regenerate, we used adult wings emerged from *sal^E/Pv^>rpr* individuals in which patterning and size defects can be scored easily. Flies were fixed in glycerol:ethanol (1:2) for 24 hours. Wings were dissected in water and then washed with ethanol. Subsequently, they were mounted in lactic acid:ethanol (6:5) and analyzed and imaged under a microscope.

The percentage of normal wings refers to normally regenerated or developed wings and was calculated according to the number of wings with a complete set of veins and interveins, as markers of normal patterning. For each sample, we scored the percentage of individuals belonging to the “normal wings” class and calculated the standard error of the sample proportion based on a binomial distribution (regenerated complete wing or not) SE =*√* p (1-p)/n, where p is the proportion of successes in the population.

### Immunochemistry

Immunostaining was performed using standard protocols. The primary antibodies used in this study were Ask1 P-Ser83 (Santa Cruz Biotechnology sc-101633 1:100, which recognizes the conserved region surrounding human P-Ser83), Ask1 P-Thr (Santa Cruz Biotechnology sc-109911 1:100, which labels the preserved region nearby mouse P-Thr845), Ptc (DSHB, 1:100), P-p38 (Cell Signalling 1:50), Mmp1 (cocktail of three antibodies: DSHB 3A6B4, 5H7B11, 3B8D12 1:100), P-Akt (S473 Cell Signalling 1:100) and P-Histone-H3 (Millipore, 1:1000). Fluorescently labeled secondary antibodies were obtained from ThermoFisher Scientific. Discs were mounted in SlowFade or ProLong (ThermoFisher Scientific) supplemented with 1 µM TO-PRO-3 (TP-3) to label nuclei. For apoptotic cell detection, we used the TUNEL assay. We employed the fluorescently labeled Alexa Fluor^®^ 647-aha-dUTP (ThermoFisher Scientific), incorporated using terminal deoxynucleotidyl transferase (Roche).

The number of mitotic cells per µm^2^ (Fig 1C) was calculated after counting the number of P-Histone-H3 (P-H3) positive cells in the anterior compartment of the wing pouch and hinge for all the genotypes shown. The number of mitosis after analyzing the stacks of confocal images was calculated using Fiji software.

### Imaginal disc culture and physical injury

Wing discs were dissected from third instar larvae in Schneider’s insect medium (Sigma-Aldrich), and a small fragment was removed with tungsten needles. To visualize Mmp1 staining, the discs were cultured in Schneider’s insect medium supplemented with 2% heat-activated fetal calf serum, 2.5% fly extract and 5 µg/ml insulin, for 5 hours at 25°C. For P-p38, discs were injured in Schneider’s insect medium and immediately fixed and stained. *Ex vivo* images were taken using a Leica SPE confocal microscope and processed with Fiji software.

### ROS scavenging

To prevent ROS production in *sal^E/Pv^>rpr* discs we pursued two different protocols. First, we inhibited ROS chemically (Fig 4A-F). To do this, standard fly food was supplemented with the antioxidant N-acetyl cysteine (NAC) (100 µg/ml) (Sigma-Aldrich). NAC treatment was dispensed on the 7^th^ day of development at 17°C. On the 8^th^ day, experimental larvae were moved to 29°C for 11 hours to promote cell death, whereas controls were transferred to a vial with standard food and moved to 29°C for the same time period. Afterward, the larvae were move back to 17°C to allow tissue recovery. Second, we decreased ROS production genetically by the ectopic expression of the Sod1 and Catalase enzymes using a recombinant fly *UAS-Sod1:UAS-Cat* (*Sod1:Cat*) in the *sal^E/Pv^>rpr* domain. For those experiments, *Sod1:Cat* and *rpr* were activated for 24 hours at 29°C

### Oxidative stress induction

Third instar larvae were transferred to vials with 5mL of special medium containing 1.3% UltraPure^TM^ LMP agarose (Invitrogen), 5% sucrose (Fluka) and the desired concentration of 0.1% H_2_O_2_(Merck) or 1ng/µl tunicamycin (Sigma-Aldrich). To avoid loss of oxidative capacity, these substances were added to the media at a temperature below 45°C. The larvae were fed for 2 hours prior to dissection and fixation of the discs. Controls without H_2_O_2_ or tunicamycin were always handled in parallel.

### Genotypes

**Figure 1.**

(A) <u>*+/+*</u> → *wUAS-rpr/+; sal^E/Pv^-Gal4/+; tubGal80^TS^/+*

<u>*Ask1^MI02915^/+*</u> → *wUAS-rpr/+; sal^E/Pv^-Gal4/+; tubGal80^TS^/Ask1^MI02915^*

<u>*Ask1^MB06487^/+*</u> → *wUAS-rpr/+; sal^E/Pv^-Gal4/+; tubGal80^TS^/Ask1^MB06487^*

(B) <u>*sal^E/Pv^>rpr*</u> → *w; ci-Gal4/LexO-rpr; sal^E/Pv^-LHG:tubGal80^TS^/UAS-GFP*

<u>*ci>Ask1^RNAi^*</u> → *w; ci-Gal4/LexO-GFP; sal^E/Pv^-LHG:tubGal80^TS^/ UAS-Ask1^RNAi^*

<u>*sal^E/Pv^>rpr ci> Ask1^RNAi^*</u> → *w; ci-Gal4/LexO-rpr; sal^E/Pv^-LHG:tubGal80^TS^/ UAS-Ask1^RNAi^*

<u>*ap> Ask1^RNAi^*</u> → *w; ap-Gal4/LexO-GFP; sal^E/Pv^-LHG:tubGal80^TS^/ UAS-Ask1^RNAi^*

<u>*sal^E/Pv^>rpr ap> Ask1^RNAi^*</u> → *w; ap-Gal4/LexO-rpr; sal^E/Pv^-LHG:tubGal80^TS^/ UAS-Ask1^RNAi^*

(C, D) <u>*sal>GFP ci>RFP*</u> → *w; ci-Gal4/LexO-GFP;*

<u>*sal^E/Pv^-LHG:tubGal80^TS^/UAS-RFP sal>GFP ci>Ask1^RNAi^*</u> → *w; ci-Gal4/LexO-GFP; sal^E/Pv^-LHG:tubGal80^TS^/ UAS-Ask1^RNAi^*

<u>*sal^E/Pv^>rpr ci>GFP*</u> → *w; ci-Gal4/LexO-rpr; sal^E/Pv^-LHG:tubGal80^TS^/UAS-GFP*

<u>*sal^E/Pv^>rpr ci> Ask1^RNAi^*</u> →*w; ci-Gal4/LexO-rpr; sal^E/Pv^-LHG:tubGal80^TS^/ UAS-Ask1^RNAi^*

**Figure 2.**

(B,C) <u>WT</u> → Canton S

(D) <u>*sal^E/Pv^>rpr*</u> → *wUAS-rpr/+; sal^E/Pv^-Gal4/+; tubGal80^TS^/+*

(E,F) <u>*sal^E/Pv^>rpr ci> Ask1^RNAi^*</u> → *w; ci-Gal4/LexO-rpr; sal^E/Pv^-LHG:tubGal80^TS^/*

*UAS-Ask1^RNAi^*

(G) <u>*hh>Ask1^WT^, GFP*</u> → *w; UAS-GFP/+; hh-Gal4/UAS-Ask1^WT^*

(H) <u>*hh>Ask1^S83A^, GFP*</u> → *w; UAS-GFP/+; hh-Gal4/UAS-Ask1^S83A^*

(I) <u>*sal>GFP ci>Ask1^WT^*</u> → *w; ci-Gal4/LexO-GFP; sal^E/Pv^-LHG:tubGal80^TS^/UAS-Ask1^WT^*

<u>*sal>GFP ci>Ask1^S83A^*</u> →*w; ci-Gal4/LexO-GFP; sal^E/Pv^-LHG:tubGal80^TS^/UAS-Ask1^S83A^*

<u>*sal^E/Pv^>rpr*</u> → *w; ci-Gal4/LexO-rpr; sal^E/Pv^-LHG:tubGal80^TS^/UAS-GFP*

<u>*sal^E/Pv^>rpr ci> Ask1^WT^*</u> → *w; ci-Gal4/LexO-rpr; sal^E/Pv^-LHG:tubGal80^TS^/UAS-Ask1^WT^*

<u>*sal^E/Pv^>rpr ci> Ask1^S83A^*</u> →*w; ci-Gal4/LexO-rpr; sal^E/Pv^-LHG:tubGal80^TS^/UAS-Ask1^S83A^*

**Figure 3.**

A) Canton S

(B) <u>*ptc>rpr*</u> → *wUAS-rpr/+; ptc-Gal4: tubGal80^TS^ /+*

(C) <u>*sal>rpr*</u> →*wUAS-rpr/+; sal-Gal4/+; tubGal80^TS^/+*

(D,E) <u>*ptc>Akt^RNAi^*</u> → *w; ptc-Gal4:tubGal80^TS^/+;UAS-AktRNAi/+*

(F,G) <u>*ptc>myrAkt*</u> → *w; ptc-Gal4:tubGal80^TS^/+; UAS-myrAkt/+*

(H-K) <u>*sal^E/Pv^>rpr hh>dp110^DN^*</u> →*w; LexO-rpr /UAS-dp110^DN^; sal^E/Pv^-LHG:tubGal80^TS^/hh-Gal4*

(L) <u>*ci>dp110^DN^*</u> → *w; ci-Gal4/UAS-dp110^DN^; sal^E/Pv^-LHG:tubGal80^TS^/+*

<u>*+/+*</u> → *w; LexO-rpr/ +; sal^E/Pv^-LHG:tubGal80^TS^/+* (control for LexO-rpr in the second chromosome) and *wUASrpr/+; sal^E/Pv^-Gal4; tubGal80^TS^* (control for mutant backgrounds)

<u>*sal^E/Pv^>rpr ci>dp110^DN^*</u> → *w; ci-Gal4/UAS-dp110^DN^; sal^E/Pv^-LHG:tubGal80^TS^/LexO-rpr*

<u>*sal^E/Pv^>rpr Akt^1/+^*</u> → *wUAS-rpr/+; sal^E/Pv^-Gal4; tubGal80^TS^; Akt^1^/+*

<u>*sal^E/Pv^>rpr Ask1^MB06487/+^ Akt^1/+^*</u> → *wUAS-rpr/+; sal^E/Pv^-Gal4; tubGal80^TS^; Akt^1^/Ask1^MB06487^*

<u>*sal^E/Pv^>rpr Ask1^MI02915/+^ Akt^1/+^*</u> →*wUAS-rpr/+; sal^E/Pv^-Gal4; tubGal80^TS^; Akt^1^/Ask1^MI02915^*

**Figure 4.**

(B,C) <u>Std food and NAC</u> →*wUAS-rpr/+; sal^E/Pv^-Gal4/+; tubGal80^TS^/+*

(E,F) <u>CTRL</u> → *w^118^; +; +*

<u>*Ask1^-/-^*</u> →*w^118^; +; Ask1^MB06487^/Ask1^MB06487^*

(G) <u>*sal>rpr, GFP*</u> → *w; sal^E/Pv^-Gal4/UAS-GFP; tubGal80^TS^/UAS-rpr*

<u>*sal>rpr, Sod1:Cat*</u> →*w; sal^E/Pv^-Gal4/UAS-Sod1:UAS-Cat; tubGal80^TS^/UAS-rpr*

**Figure 5.**

(A,C) <u>*sal^E/Pv^>rpr*</u> → *w; LexO-rpr/ +; sal^E/Pv^-LHG:tubGal80^TS^/+*

(B,D) <u>*sal^E/Pv^>rpr ci>Ask1^RNAi^*</u> → *w; ci-Gal4/LexO-rpr; sal^E/Pv^-LHG:tubGal80^TS^/UAS-Ask1^RNAi^*

(E-H) <u>*en>Ask1^RNAi^, GFP*</u> →*w; en-Gal4/UAS-GFP; UAS-Ask1^RNAi^/+*

**S1 Figure.**

(A)*w^118^; +; +*

(B) <u>*Ask1^+^/Ask1^+^*</u> → *w^118^; +; +*

<u>*Ask1^MB06487^/Ask1^MB0647^*</u> → *w^118^; +; Ask1^MB06487^/Ask1^MB06487^*

<u>*Ask1^MB06487^/Def(3R)BSC636*</u> → *w^118^; +; Ask1^MB06487^/Def(3R)BSC636*

(C) <u>*hh>Ask1^RNAi^,GFP*</u> → *w; UAS-GFP/+; hh-Gal4/UAS-Ask1^RNAi^*

**S2 Figure.**

(A) wt and *ptc>rpr* → *wUAS-rpr/+; ptc-Gal4: tubGal80^TS^ /+*

<u>*sal^E/Pv^>hid*</u> → *w; sal^E/Pv^-Gal4/+; tubGal80^TS^/UAS-hid*

<u>wt and physically injured</u> → Canton S

**S3 Figure.**

**(A-D)** <u>*ci>dp110^DN^*</u> → *w; ci-Gal4/UAS-dp110^DN^; sal^E/Pv^-LHG:tubGal80^TS^/ LexO-rpr*

## Acknowledgments

We thank Josep F. Abril, Irene Martínez and Irene Pardo for their help during the initial stages of this work. We thank Hugo Stocker for helpful discussions and providing reagents. We also thank Marta Morey, Haritz Plazaola, Qui Zhu, Alejandra Fernández and Elena Vizcaya for discussions. Stocks obtained from the Bloomington Drosophila Stock Center (NIH P40OD018537) were used in this study. We thank the Genomics Resource Center, supported by NIH grant 2P40OD010949, for the Ask1 clone. We also thank Manel Bosch at the confocal facility of the Scientific and Technological Centers (CCiT) of the University of Barcelona for his help. This project was funded by the Spanish Ministerio de Economía y Competitividad, Spain grant BFU2015-67623-P to FS and MC and by grant BFU2016-77587-P to M.M.’s lab.

## Supporting Information Legends

**S1 Figure.** (A) Mitotic (P-H3 positive) cells co-localize with high levels of Ask1 P-Thr. Lower right images: Zoom of mitotic cells from the white square. TP-3: TO-PRO-3 nuclear staining. (B) The basal levels of P-Thr in wild-type (WT) discs (mitotic and interphase cells) decrease in the *Ask1^MB06487^* homozygous mutant background and are strongly reduced in the *Ask1^MB06487^* mutant background over the deficiency *Def(3R)50636.* Quantification of mean pixel intensity for P-Thr in mitotic and for P-Thr in non-mitotic (basal levels) cells are shown in separate graphics. P-Thr mean Pixel intensity in mitotic cells of the wild-type discs (+/+), 113.92±20.60, S.D. (n=10); of the *Ask1^MB06487^/Ask1^MB06487^* discs, 104.43±9.54, S.D. (n=10); and of the *Ask1^MB06487^/Def(3R)BSC636* 88.37±13.03, S.D. (n=10). Mean pixel intensity of the basal levels in interphase cells of the wild-type, 50.44±6.27, S.D. (n=20); of the *Ask1^MB06487^/Ask1^MB06487^*, 41.65±4.80, S.D. (n=10); and of the *Ask1^MB06487^/Def(3R)BSC636*, 27.22±5.73, S.D. (n=18). (C) Inhibition of the endogenous Ask1 P-Ser83 levels after expressing the *UAS-Ask1^RNAi^* in the posterior compartment, using the *hh-Gal4* driver. Scale bars: 50µm.

**S2 Figure.** (A) Left: wild-type disc stained with P-Ser83. Right: disc in which apoptosis has been induced with the driver *ptc-Gal4 UAS-rpr* (*ptc>rpr*) and stained with P-Ser83 (n=5) showing extensive increase in P-Ser83 in the undamaged cells. (B) S*al^E/Pv^>hid* discs stained with P-Thr (n=12) and P-Ser83 (n=5). TP-3: TO-PRO-3 nuclear staining. Purple: *sal^E/Pv^>rpr* ablated zone. (C) After physical injury, Ask1 P-Ser83 (n=6) accumulates in cells at the wound edges. Square: magnified image. Scale bars: 50µm.

**S3 Figure.** (A) Design of the experiments in B,C and D with *sal^E/Pv^>rpr* cell death (purple) and blocking Pi3K (*ci>dp110^DN^*) in the anterior compartment (black). (B-D) Staining of P-Akt (B), Ask1 P-Ser83 (C) and Ask1 P-Thr (D). Note that P-Akt and PSer83 localization is reduced but not P-Thr. Dotted white line indicates anterior-posterior boundary. TP3: TO-PRO-3 nuclear staining. Scale bars: 50µm.

